# A first-takes-all model of centriole copy number control

**DOI:** 10.1101/2020.09.23.309567

**Authors:** Marco António Dias Louro, Mónica Bettencourt-Dias, Jorge Carneiro

## Abstract

How cells control the numbers of its subcellular components is a fundamental question in biology. Given that biosynthetic processes are fundamentally stochastic it is utterly puzzling that some structures display no copy number variation within a cell population. Centriole biogenesis, with each centriole being duplicated once and only once per cell cycle, stands out due to its remarkable fidelity. This is a highly controlled process, which depends on low-abundance rate-limiting factors. How can exactly one centriole copy be produced given the natural variation in the concentration of these key players? Hitherto, tentative explanations of this control evoked lateral inhibition-or phase separation-like mechanisms emerging from the dynamics of these rate-limiting factors, but how centriole number is regulated remains unclear. Here, we propose a novel solution to centriole copy number control based on the assembly of a centriolar scaffold, the cartwheel. We hypothesise that once the first cartwheel is formed it continues to elongate by stacking the intermediate cartwheel building blocks that would otherwise form supernumerary structures. Using probability theory and computer simulations, we show that this mechanism may ensure formation of one and only one cartwheel over a wide range of parameter values at physiologically relevant conditions. By comparison to alternative models, we conclude that the key signatures of this novel mechanism are an increasing assembly time with cartwheel numbers and that stochasticity in cartwheel building blocks should be converted into variation in cartwheel numbers or length, both of which can be tested experimentally.

**Author summary:** Centriole duplication stands out as a biosynthetic process of exquisite fidelity in the noisy world of the cell. Each centriole duplicates exactly once per cell cycle, such that the number of centrioles per cell shows no variance across cells, in contrast with most cellular components that show broadly distributed copy numbers. We propose a new solution to the number control problem. We show that elongation of the first cartwheel, a core centriolar structure, may sequester the building blocks necessary to assemble supernumerary centrioles. As a corollary, the variation in regulatory kinases and cartwheel components across the cell population is predicted to translate into cartwheel length variation instead of copy number variation, which is an intriguing overlooked possibility.

## Introduction

Cellular biosynthetic processes are inherently stochastic due to key intervening molecules that exist in low abundance. Importantly, variation in the concentrations of specific proteins is systematically observed over time in single cells [1] and within populations of genetically identical cells in the same state [2]. In this context, it is paradoxical that some cellular components have little or zero variance in copy number across a cell population. This suggests an underlying biosynthetic process of exquisite fidelity, whose deterministic-like control mechanism demands explanation.

Centriole duplication stands out as a paradigm of an extraordinary number control mechanism [3, 4]. Typical proliferating vertebrate cells contain two centrioles in G1. During S-phase, these centrioles are duplicated once and only once. After mitosis, each daughter cell inherits a single pair of centrioles. Centriole over-or underproduction generally leads to mitotic defects and cell-cycle arrest or cell death. [5–7]; thus, rarely observed in healthy tissues.

In human cells, centriole numbers are depend on the levels of key proteins, namely Plk4, STIL, and SAS-6. Depletion of either one efficiently abrogates centriole duplication and their overexpression triggers the production of random supernumerary centrioles in a dose-dependent manner [8–11]. Quantitative proteomics revealed that Plk4, STIL, and SAS-6 are relatively low-abundance proteins at physiological conditions [12] although centriole duplication proceeds efficiently at a relatively broad range of their concentrations [13, 14]. Although active degradation of Plk4, STIL, and SAS-6 via the cell cycle machinery and/or their own activity [15–17] may allow their levels to be kept within a given range, quasi-deterministic centriole duplication requires mechanisms that buffer stochasticity in Plk4, STIL, and SAS-6 levels.

Several studies have proposed that these proteins can enact spatial control of centriole biogenesis by cooperatively focusing at a single site [18, 19], facilitating local centriole assembly and inhibiting it elsewhere. Recently published mathematical models predict these dynamics based on different mechanisms [20–22]. In principle, these models allow for robust centriole duplication by restricting its key regulators in space. However, they do not address the question of how centriole number control is directly achieved at a molecular level, especially concerning how local Plk4, STIL, and SAS-6 levels are converted into discrete numbers of centrioles.

We propose that centriole number control can be achieved by controlling the numbers of a core structure called the cartwheel. The cartwheel is assembled following Plk4, STIL, and SAS-6 focusing and acts as a platform around which one and only one centriole is built. The central hub of the cartwheel consists of a multi-layered stack, where each ring-like layer is composed of nine SAS-6 homodimers [23–25]. Ring formation via oligomerisation and stacking depend on structural properties of SAS-6, at least partly [26, 27]. We hypothesise that growing SAS-6 oligomers can be stacked up on the first-formed cartwheel hub, heretofore simply referred to as cartwheel, thus preventing the formation of supernumerary structures.

Here, we assess this hypothesis by modelling stochastic cartwheel assembly. Our model can be summarized as the competition between two processes: combinatorial multi-step oligomerisation leading to new cartwheels, and sequestering of intermediate oligomers by previously assembled cartwheels, resulting in new rings being added to the stack. We show that elongation of a single cartwheel, by stacking, can efficiently prevent supernumerary cartwheels over a wide range of model parameters and in distinct theoretical scenarios. Finally, we compare our model to a general mechanism of structure number control by negative feedback and find a set of specific predictions: first, cartwheel assembly time is increasingly delayed with the formation of each extra cartwheel; and second, both cartwheel numbers and length may vary as a function of SAS-6 levels.

## Results

### Sequestering of SAS-6 intermediates inhibits formation of supernumerary cartwheels

We explore a simple principle of biosynthetic regulation, where the the final product of a multi-step assembly pathway inhibits the synthesis of further products by incorporating intermediate molecules as they form. As per our hypothesis, this boils down to a competition between assembly of new cartwheels by oligomerisation of SAS-6 dimers and elongation of the first-formed cartwheel by stacking these oligomers.

We first illustrate this principle by calculating the fate of a SAS-6 dimer that may either integrate the first-formed cartwheel or give rise to an extra one. We consider a focal volume around a pre-existing centriole that is large enough to allow for mass action dynamics. We assume that SAS-6 oligomers *C*_*i*_, containing 1 ≤ *i<r* dimers, can bind a free dimer irreversibly to form *C*_*i*+1_ oligomers, with constant rate *k*_*on*_ – the “oligomerisation” rate. Upon reaching *C*_*r*_, we consider that an independent cartwheel ring has formed. The completion of this new cartwheel ring is the minimal condition the establishment of a distinct centriole. Alternatively, at each of the *C*_*i*_ intermediate steps, the oligomers can be sequestered by binding to the previously formed cartwheel also with a constant rate *k*_*s*_ – the “stacking” rate.

Consider a given SAS-6 dimer (*C*_1_). Given enough time this dimer will either be sequestered by the elongating cartwheel or it will progress down the assembly path to integrate a new cartwheel ring, which we postulate to yield an extra centriole. As time approaches infinity, the probability that the dimer is incorporated into to a new independent cartwheel ring *C*_*r*_ tends asymptotically to a maximum given by (the derivation of Eq. 1 is provided in Materials and Methods):

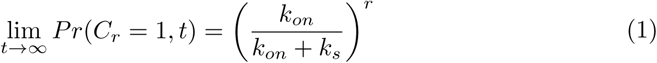

The right-hand side of the equation represents the probability that the original dimer undergoes *r −* 1 consecutive oligomerisation steps without any stacking reaction event, thus yielding a new complete cartwheel ring. This probability Eq. 1 decreases exponentially with ring size *r* and as a power-law function of the stacking rate *k*_*s*_ (Fig. 1). In nature, cartwheels display a highly conserved nine-fold symmetry (*r* = 9), although rings of different sizes have been observed *in vitro* [28]. In these conditions, when *k*_*on*_ = *k*_*s*_, i.e. when oligomerisation and stacking rates are identical, the probability of forming supernumerary cartwheels is 0.2%. Interestingly, even if the value of *k*_*s*_ is half that of *k*_*on*_, this values rises only to 2.6%. Thus, under this simple model, efficient cartwheel number control does not require relatively higher stacking rates. We emphasise that this is possible due to the number of intermediate steps before a new cartwheel ring is assembled.

**Fig 1.**
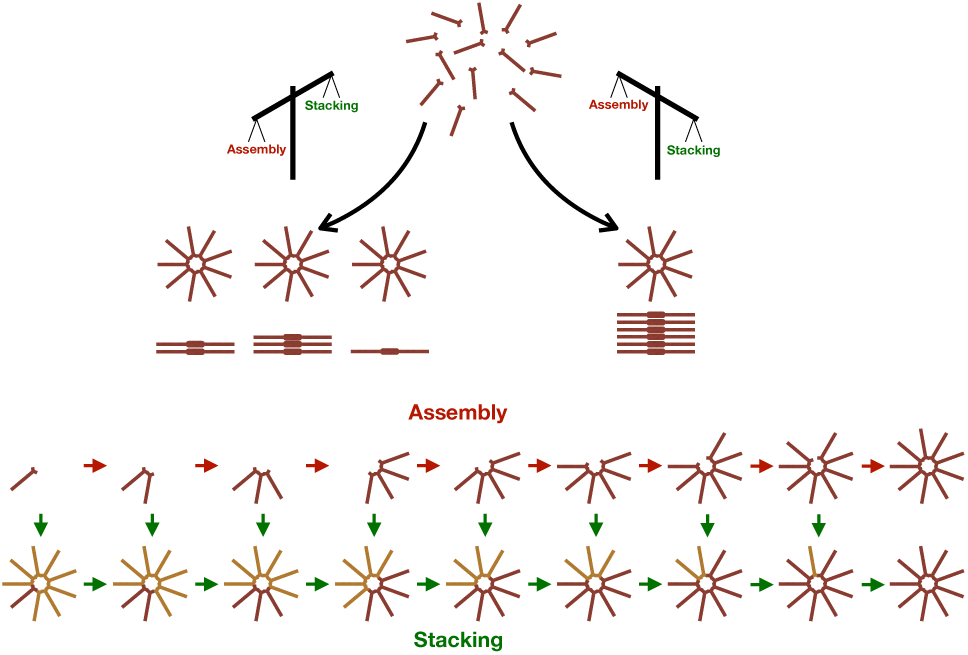
General scheme of the model. The SAS-6 dimer is the basic subunit of the carwheel. Higher-order oligomers can arise through oligomerisation of SAS-6 dimers. Once oligomers grow to assemble a complete ring, it is assumed that an independent cartwheel is formed. After the first cartwheel is formed it can elongate by stacking intermediate oligomers onto the top ring layer. Thus, stacking can deplete the pool of available dimers and oligomers necessary to assemble supernumerary cartwheels.

**Fig 2.**
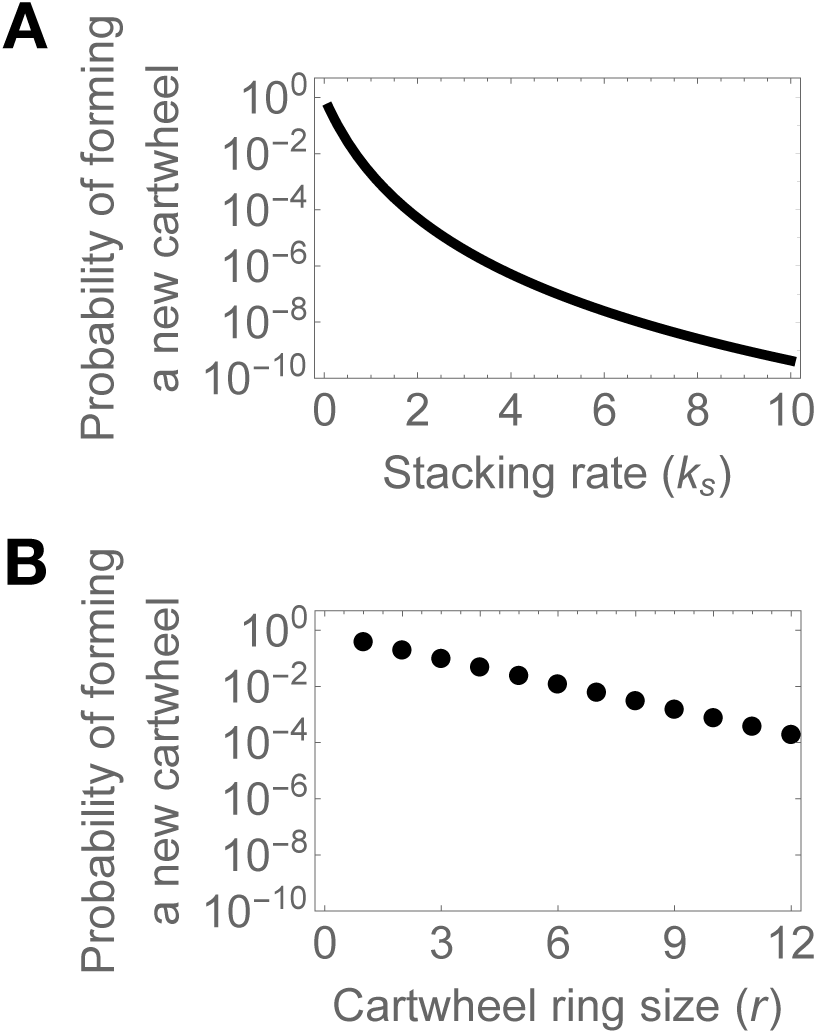
The probability of forming extra cartwheels depends negatively on the stacking rate and ring size. Equation Eq. 1 was evaluated numerically at the indicated parameter values. We set *k*_*on*_ = 1.

Despite its analytic insights, the previous analysis is an oversimplification of the biochemical processes involved in cartwheel assembly. Importantly, ring formation has been described as a combinatorial oligomerisation process, where intermediates of different sizes can combine into higher-order structures [27, 29]. For example, two molecules containing four (*C*_4_) and five dimers (*C*_5_) can combine, skipping over multiple steps in the assembly pathway, to form a complete ring. This also implies considering an entire molecular population, rather than tracking the fate of a single molecule participating in successive first-order reaction events.

To explore these potential issues, we formulated a more realistic model of cartwheel assembly. We assume that SAS-6 expression, degradation, and recruitment to the focal volume, result in a constant net influx of *C*_1_ dimers into the focal volume, at a rate σ. This is based on fluorescence measurements of SAS-6 intensities indicate that it accumulates during S-phase, peaking in G2 [12, 30]. *C*_*i*_ and *C*_*j*_ molecules can interact to produce higher-order *C*_*i*+*j ≤r*_ oligomers, at a constant rate *k*_*on*_. In turn, *C*_*i*+*j<r*_ can dissociate into *C*_*i*_ and *C*_*j*_ at a constant rate *k*_*off*_.

We introduce the 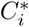 notation to represent stacks containing at least one layer of complete rings containing *r* = 9 dimers, topped by an incomplete ring containing 0 ≤ *i<r* oligomers. Note that 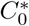 substitutes *C*_*r*_. We postulate the existence of a stacking mechanism analogous to ring assembly by oligomerisation. When a 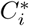 stack binds a freely-diffusing *C*_*j*_ oligomer it forms a 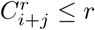 stack. This can be visualised as a ring being assembled on top of a stack, aligned with the layers underneath [25]. We assume that all oligomerisation reactions yielding complete cartwheel rings as well as all stacking reactions are irreversible.

The detailed model depicts cartwheel assembly as an oligomerisation and stacking process, under a constant influx of SAS-6 dimers. This model allows for the assembly of multiple cartwheels and their elongation. In fact, the number of SAS-6 dimers in the focal volume increases monotonically as a function of time. Thus, it does not reach a steady-state, making it quite cumbersome for deriving analytical solutions. Readers may refer to Materials and Methods for detailed formulation of the model.

We resorted to numerical simulations to study the behaviour of this model. We set a fixed time window in which a cell is permissive to centriole biogenesis, roughly corresponding to S-phase, in which SAS-6 is shuttled into the centrosome. Thus, neither *C*_*i*_ intermediates nor cartwheels are present at the beginning of the simulations. In addition, this poses a challenge to our hypothesis as stacking may not be efficient enough to prevent other large oligomers (e.g. *C*_*i*_) from seeding extra cartwheels.

Figure 3 shows the temporal dynamics of cartwheel formation and numbers of *C*_*i*_ intermediates. In the absence of stacking (Fig.3A,D), supernumerary cartwheels rapidly appear and keep forming, on average, at an approximately linear rate. When all four parameters are set to the same value (Fig. 3B,E), the emergence of the first cartwheel coincides with the depletion of *C*_*i*_ intermediates, and supernumerary cartwheels are formed only in 2.3% of the simulations. When σ is increased with respect to the other parameters (Fig.3C,F), mimicking, for example, SAS-6 overexpression, supernumerary cartwheels appear at a decelerating rate, while reducing the levels of building blocks. See also Fig. S1 Concomitantly, assembled cartwheels elongate continuously (Fig. 4, Fig. S2). For reference, note that in the absence of stacking, cartwheels attain a maximum length of one ring. When stacking is enabled, building blocks are equally distributed by existing cartwheels. This can be observed in “overexpression” conditions (Fig. 4C plot). At first, when the first cartwheel is present, its elongation rate is maximal. When the second cartwheel is formed in all simulations, and formation of additional cartwheels becomes less favourable, the average elongation rate for both cartwheels equilibrates. The difference in assembly times dictates that the first cartwheel is longer, on average, than the second cartwheel, at the end of the simulations. Notwithstanding, inherent stochasticity in molecular influx and stacking result in a time-dependent increase in cartwheel length variability.

**Fig 3.**
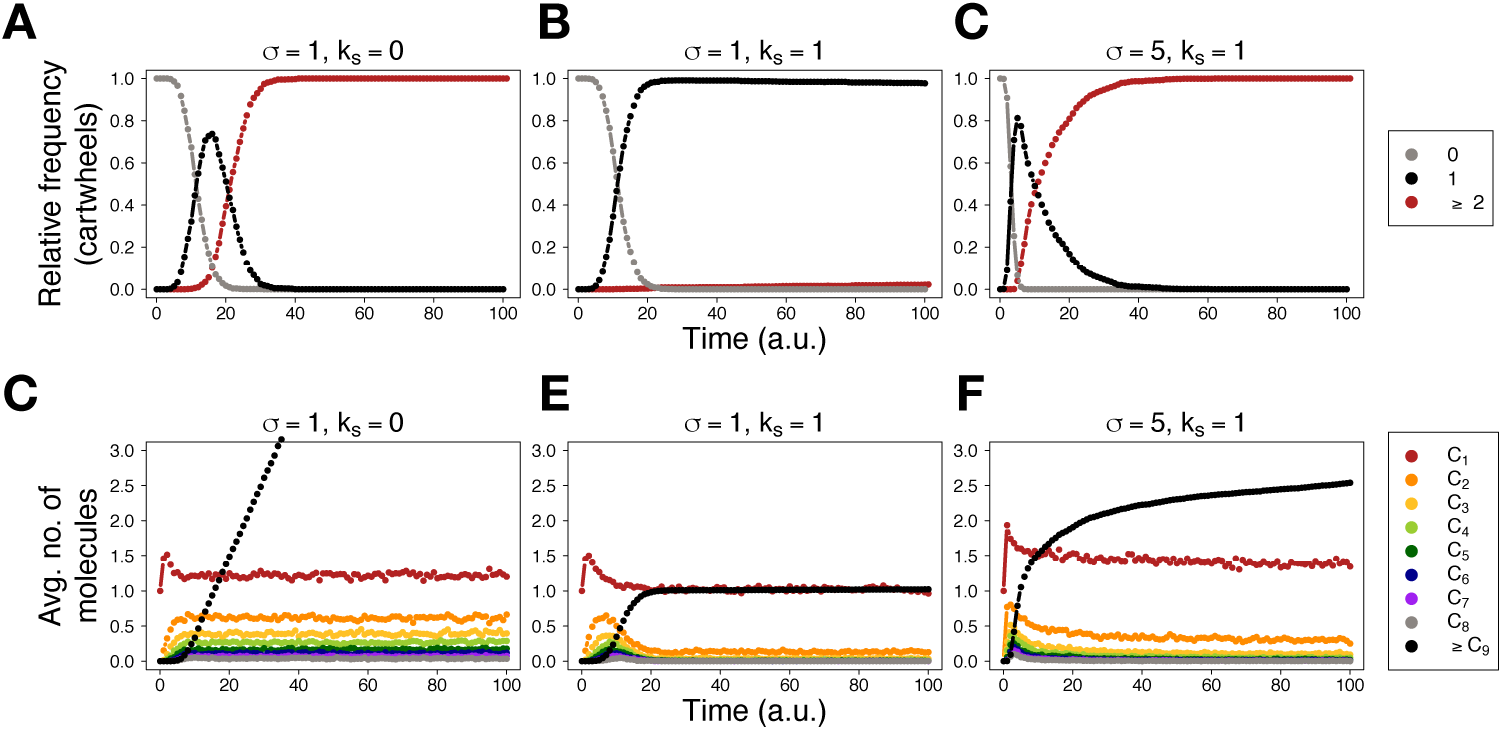
Stacking causes depletion of intermediates and inhibits the formation of supernumerary cartwheels. (A-C) - relative frequency of simulations containing 0 (grey), 1 (black), and 2 or more (red) cartwheels as a function of time, for the indicated values of σ and *k*_*s*_. (D-F) - average number of molecules across all simulations, as a function of time, for the indicated values of σ and *k*_*s*_. We performed 1000 simulations with no SAS-6 molecules initially present and stopped them at time *t* = 100; time *t* is measured in arbitrary units (a.u.). For all simulations, *k*_*on*_ = *k*_*off*_ = 1. Note that *k*_*s*_ = 0 represents absence of stacking. See also Methods section for default initial and stopping conditions, and parameter values for the simulations.

**Fig 4.**
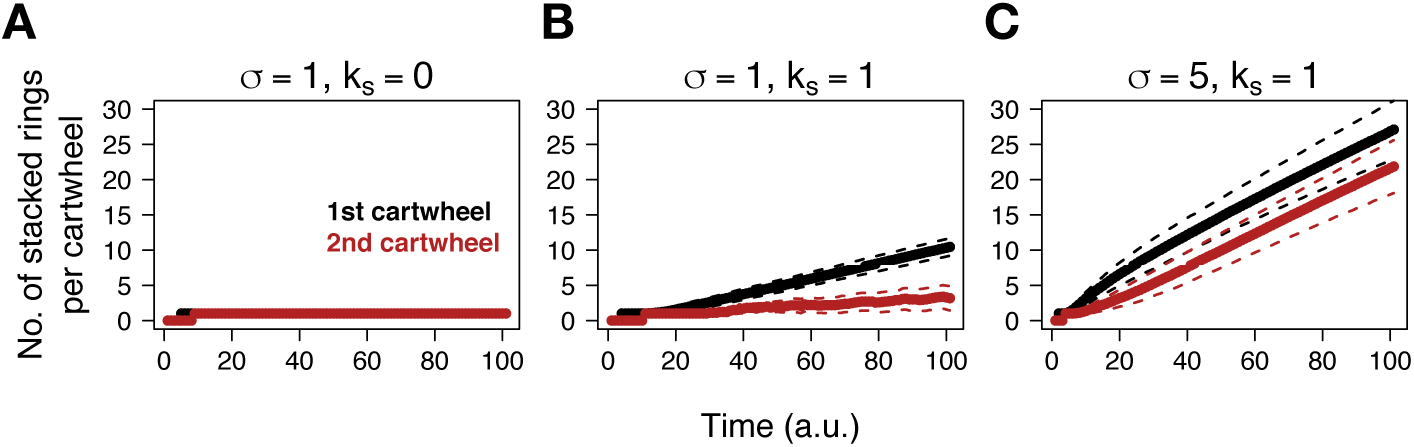
Cartwheels elongate continuously and show a distribution of lengths along time. The average number of complete rings present in the first (black) and second (red) cartwheels across all simulations are plotted as a function of time, for the indicated values of σ and *k*_*s*_. The dashed lines indicate the average *±* standard deviation. We used default simulations conditions as indicated in Figure 3 and in the Methods section. Note that *k*_*s*_ = 0 represents absence of stacking.

In qualitative terms, we obtain similar results as in the previous model. First, stacking can efficiently suppress the formation of supernumerary cartwheels without the need of high stacking rates. Second, we observe a similar dependence on ring size on the probability of forming supernumerary cartwheels (Fig. S3).

### Formation of one and only one cartwheel is achieved with high fidelity for a wide range of parameter values

Up to this point, we have established a simple principle of inhibition-by-stacking. The postulated mechanism can ensure formation of one and only one cartwheel with high fidelity in unbiased conditions in which stacking does not occur at high rates. However, it is pertinent to explore a wider region of the parameter space and evaluate model outcomes, to identify the conditions guaranteeing proper cartwheel number control.

We conducted simulations of the model for pairwise combinations of parameter values in [0.1,2.0], whilst keeping other parameters at the value of 1 (Fig. 5A, see also Fig. S4). Specifically, we are interested in characterizing the relationship between the influx parameter σ, that ultimately limits maximum cartwheel number and length, and the other reaction parameters. In the designated range, there will be, on average, between 10 and 200 SAS-6 dimers at the end of the simulations. This means that 1-22 complete rings may form, on average. It has been estimated by quantitative proteomics that 138 dimers of SAS-6, on average, are present near centrioles after cartwheel formation, potentially yielding cartwheels of up to 15 complete rings in length [12]. The dimensions of the SAS-6 focus near centrioles also support a cartwheel length of 7-17 rings [30]. Therefore, our analysis covers a range of molecular abundances that can be related to strong SAS-6 depletion through to mild overexpression.

**Fig 5.**
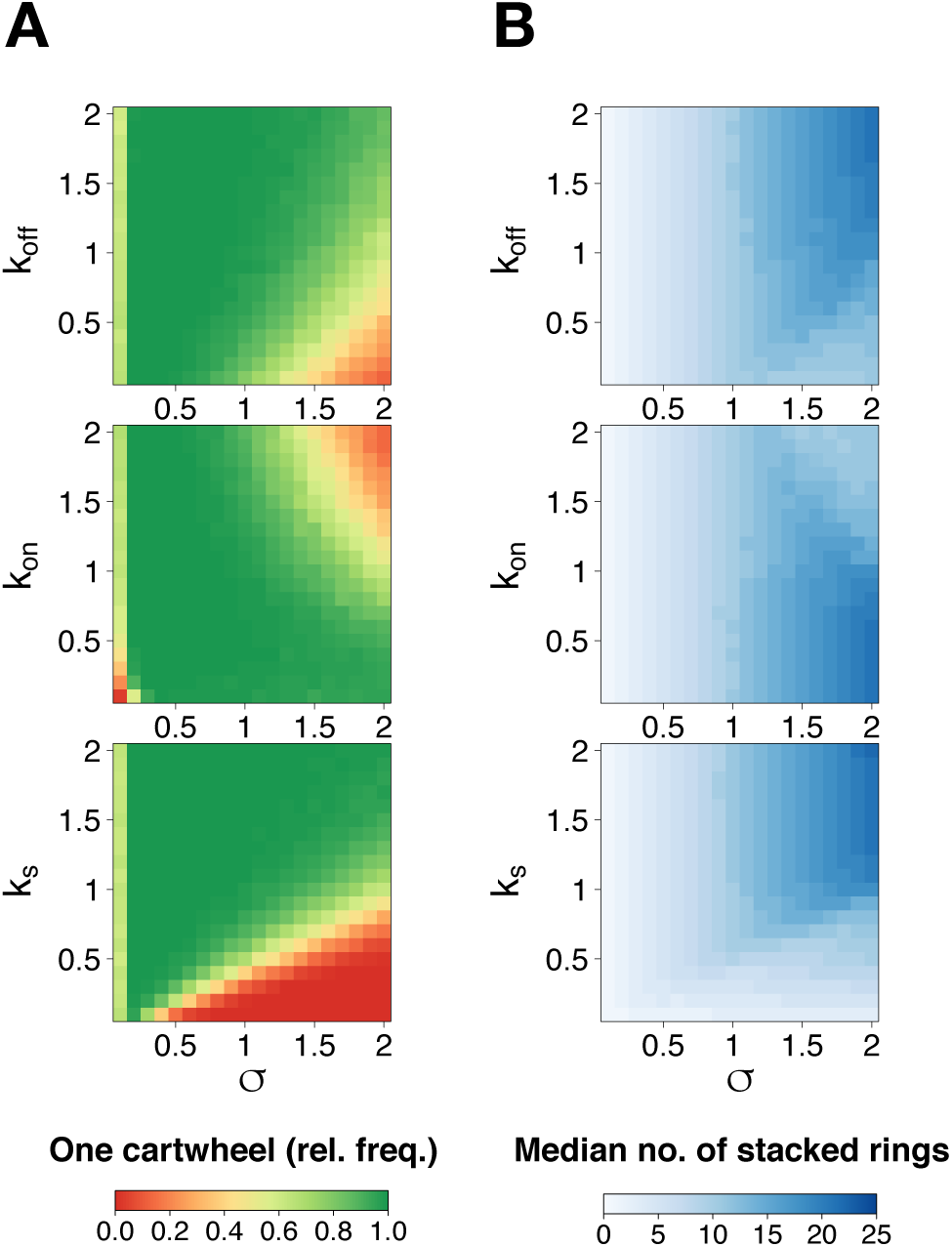
Probability of forming one and only one cartwheel as a function of influx and reaction parameters. Relative frequency of simulations containing a single cartwheel at time *t* = 100, under the model assuming irreversible ring assembly and stacking. We varied the indicated parameter values in steps of 0.1, yielding a total of 400 pairwise combinations of parameter values. We used default simulations conditions as indicated in Figure 3 and in the Materials and Methods section.

As can be expected, formation of extra cartwheels is favoured for high *k*_*on*_/low *k*_*off*_ and/or high values of σ and inhibited for low *k*_*on*_/high *k*_*off*_ and/or low σ. In addition, for low values of and σ, the formation of even the first cartwheel is almost disabled for concomitantly low *k*_*on*_ but seemingly insensitive to changes in *k*_*off*_.

Nevertheless, cartwheel formation shows greater sensitivity to changes in *k*_*s*_ in comparison to *k*_*on*_ and *k*_*off*_, for the considered parameter values. As previously observed, the formation of supernumerary cartwheels is promoted for low *k*_*s*_ and hindered for high *k*_*s*_, and decreases sharply as σ increases.

With respect cartwheel elongation, we note once again an opposite trend to cartwheel formation – if fewer cartwheels form, existing ones elongate at a higher rate (Fig. 5B, see also Fig. S5). Therefore, we obtained higher median cartwheel lengths for high σ together with low *k*_*on*_, high *k*_*off*_ and high *k*_*s*_.

Overall, a single cartwheel is formed in the majority of parameter combinations and for a wide range of SAS-6 levels. In addition, if SAS-6 influx is too low, cartwheel assembly is likely to fail. On the other hand, if it is too high, supernumerary cartwheels become more frequent. These results are similar to the effects of SAS-6 depletion or overexpression on centriole duplication, respectively.

Irreversible cartwheel assembly and stacking is a simplifying assumption that facilitates the definition of what constitutes a new and definitive structure, since cartwheels cannot disassemble. While we reasoned that neither process would be biased in this way, it is possible that the assumption of irreversible ring-forming and stacking reactions contributes to robust cartwheel number control.

Thus, we modified the model assuming that complete rings can also break, at a rate *k*_*off*_, and the top most layer of a stack can dissociate, with a rate *k*_*u*_, (see Materials and Methods) and asked the probability of forming one and only one cartwheel vary as a function of model parameters. We set *k*_*on*_ = *k*_*off*_ = *k*_*u*_ = 1 such that ring assembly and disassembly, as well as stacking and “unstacking”, occur at equal rates for all combinations of parameter. We tested *k*_*s*_ = 1 and *k*_*s*_ = 5.

Under identical conditions as before, we observed that the probability of forming one and only one cartwheel is lower when ring formation and stacking are reversible (Fig. S6). Interestingly, our simulations showed that the first cartwheel is more “stable”, whereas the second one undergoes frequent assembly disassembly. For example, it appears in 93% of the simulations at some point but is only present at the end of 28%, for *k*_*s*_ = 1 (Fig. S7A). Formation of one and only one cartwheel can still be achieved by increasing *k*_*s*_ (Fig. S7B). Therefore, since these results are not qualitatively different from our previous analysis, we conclude that irreversible ring formation and stacking facilitate but are not strictly necessary for robust cartwheel number control.

Altogether, our analysis reveals that the postulated stacking mechanism allows for cartwheel numbers to be strictly controlled within a broad region of the parameter space, and is robust to stochastic fluctuations in SAS-6 availability. The model can be summarized as follows: at first, SAS-6 are produced and combine until forming a complete rings which seeds a new cartwheel. Afterwards, the building blocks can be partaken by the existing cartwheel(s) or give rise to new ones. For the same influx rate, if there are fewer cartwheels, they elongate faster and are likely longer. If there are more cartwheels, they elongate at a slower pace and are likely shorter. Conversely, it is more likely that extra cartwheels are produced if there are fewer cartwheels than if there are more. In other words, the assembly time of additional cartwheels increases with the number of existing cartwheels. In practice, there is an implicit negative feedback on cartwheel assembly as existing cartwheels “consume” intermediates via stacking. Finally, due to stochastic influx of SAS-6 and reaction dynamics, the model predicts variance in both cartwheel numbers and length.

### Comparison between different scenarios for structure number control

Up to now, we have quantified the efficiency of a hypothetical mechanism of number control and its consequences. Since, how it is currently unknown how cartwheel stacks form, we decided to explore alternative scenarios to better understand the limits of our model and how it could be experimentally verified.

First, a key assumption of the stacking model is that cartwheels can elongate indefinitely. However, it is likely that there are biophysical constraints to cartwheel length. So, we considered a variant of the stacking model where the cartwheel can elongate up to a maximum length. We now assume that cartwheels can contain at most *h* rings. The stacking reaction onto a specific cartwheel stops when it reaches this maximum length. This variant represents a general scenario where there is some extrinsic or intrinsic upper limit to cartwheel length. The two variants will heretofore be referred to as “unlimited” and “limited” stacking models, respectively.

Second, it has been proposed that the number of subcellular structures can be regulated by self-inhibitory signals [4]. We postulated that cartwheel assembly triggers a negative feedback loop that blocks assembly of additional cartwheels. We assume d that the feedback pathway is activated in a gamma-distributed time after the first cartwheel formation event. The use of a gamma-distribution with a shape parameter α and rate parameter ρ reflects a linear signalling cascade with α components, whose residence times are exponentially distributed, at the end of which cartwheel assembly is arrested. The feedback model is to be interpreted as a generic model for structure number control and not as a specific molecular pathway at play during centriole biogenesis.

In a nutshell, the feedback model represents an alternative scenario for structure number control, whereas the limited stacking model features a pertinent alternative to one of the main assumptions of the original model. The main purpose here is to compare the behaviour of these different models with the unlimited stacking model. We performed 1000 simulations of the three models using the default settings (σ = *k*_*on*_ = *k*_*off*_ = *k*_*s*_). In addition, we simulated the models without stacking (*k*_*s*_ = 0) and feedback (α = ρ = 0). For the feedback model, we set α = ρ = 1. This parameterisation yields an exponential distribution with mean 1. In effect, it represents a feedback mechanism enacted by a single component and the best case scenario for structure number control, as feedback is likelier to be almost instantaneous. To clearly separate the unlimited and limited stacking models, we set *h* = 6 when simulating the latter, which should allow for the first cartwheel to attain full length before the simulation terminates.

As cartwheels cannot elongate in the absence of stacking, we reasoned the most general readout to compare between the different models is the distribution of cartwheel assembly times (Fig. 6). In the absence of both stacking and feedback, as previously observed, the second cartwheel appears quickly after the first one in all simulations. We observed that the assembly times of the first and the second cartwheels were similarly distributed. In the limited stacking model, the second cartwheel was formed in 100% of the simulations. As in the unlimited stacking model, there is a time-window during which supernumerary cartwheels scarcely assemble, after which they appear across all simulations following an approximately normal distribution. Indeed, this latter period corresponds to the time when the first cartwheel tends to be fully elongated. By increasing *h*, the distribution of second cartwheel assembly times is shifted towards higher times (Fig. S8A). Thus, if *h* is high enough, the unlimited and limited stacking models will be indistinguishable in a fixed time-window. In this case, cartwheel assembly times are dispersed throughout the whole time-window for cartwheel assembly. In the feedback model, there is an exponentially-distributed time window of opportunity for the appearance of extra cartwheels. Thus, we observed that the second cartwheel is formed relatively early (at *t* ≤ 30) and rarely (2.2% of the simulations). See Figs. S8B and S8C for additional examples.

**Fig 6.**
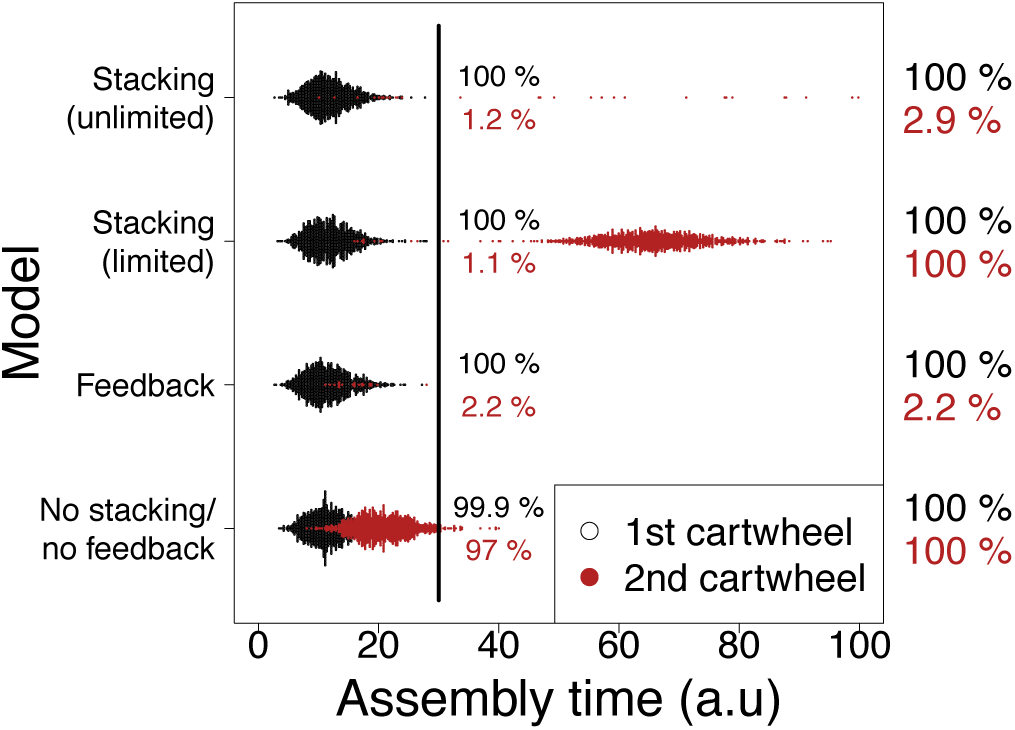
Assembly times of the first and second cartwheels under different models. The dots indicate the assembly times of the first (black) and second (red) cartwheels in 1000 simulations of the indicated models. The percentages of first (black) and second (red) cartwheels present at *t* = 30 and when the simulation terminates highlight that supernumerary cartwheels appear earlier in the feedback model when compared to the stacking model(s). We used default simulations conditions as indicated in Figure 3 and in the Materials and Methods section. For the stacking (limited and unlimited) models *k*_*s*_ = 1. Otherwise, *k*_*s*_ = 0. In the limited stacking model we set *h* = 6. In the feedback model, we set *ρ* = *α* = 1

In conclusion, the limited stacking model allows for cartwheel number control if *h* is not reached within the time-window of constant SAS-6 influx. Under these conditions, it is indistinguishable from the unlimited stacking model. Otherwise, supernumerary cartwheels are likely to appear. Note that in this case, cartwheel length may still show variation if one or more stacks of sub-maximal length are built within the permissive time window. Concerning the feedback model, extra cartwheels can only be assembled rapidly after the first one. This marks a difference from the stacking models, under which we expect the assembly time of supernumerary cartwheels to be delayed with respect to the first one.

## Discussion

In this article, we hypothesised that centriole number control can be exerted at the level of cartwheel assembly. We postulated that there is competition between forming new cartwheels and elongating existing ones for a common pool of SAS-6. Using stochastic models, which we solve analytically and with simulations, we showed that if the first cartwheel that forms is able to elongate and stack up excess SAS-6 intermediates, this can efficiently prevent formation of extra cartwheels and enable faithful centriole duplication. We showed that the efficiency depends on two fundamental aspects: (1) the length of the sequence of intermediates at which cartwheel assembly can be interrupted by stacking, and (2) the relationship between the SAS-6 influx rate and the stacking rate. For typical nine-fold symmetrical cartwheels, our results indicate that number control can be guaranteed for a wide range of model parameter values. Furthermore, the model makes the non-trivial prediction that cartwheel assembly times should increase with cartwheel numbers and that cartwheel length should be distributed in a cell population, reflecting the variance in SAS-6 influx. These two properties distinguish our model from other commonly cited number control mechanisms.

### Model predictions and limitations

Our model predicts that successive cartwheel formation events are increasingly delayed. Existing cartwheels elongate at some constant rate until a new one is formed. Since the model is inherently stochastic, both cartwheel numbers and length are distributed and their probability distribution changes in time.

The number of cartwheels increases as a function of the SAS-6 dimer influx rate in our model in accordance with what is experimentally observed upon SAS-6 depletion or overexpression. However, the influx parameter represents in a single average quantity a complex process that also includes recruitment via Plk4 and STIL, among others, which is known to be spatially regulated [18, 19], as well as post-translational degradation [15–17]. Thus, there may be a nonlinear relationship between SAS-6 levels and the influx parameter. On a different note, there is evidence suggesting that SAS-6 oligomers are relatively unstable [31], with simulations showing that SAS-6 alone cannot form cartwheels efficiently [29]. Our model is not necessarily incoherent with these studies. First, the measurements of oligomerisation kinetics were performed using truncated SAS-6 N-termini, which may differ from those in the full protein. Second, other factors may facilitate SAS-6 oligomerisation *in vivo*. If these factors are not limiting with respect to SAS-6, then the approximations in our model may still hold.

Concerning cartwheel length, there is direct and indirect evidence that it may be variable. It is well accepted that average cartwheel length is rather conserved within but may differ between distinct cell types and organisms [32]. Whereas recent studies found cartwheel length variation [26, 33], it is unclear if this represents changes in the number of stacked rings. Notwithstanding, cartwheels undergo length fluctuations during the cell cycle in *Chlamydomonas* and *Spermatozopsis* [34, 35] and they disappear during mitosis in human cells (reviewed in [24, 25]). Indeed, measurement of the SAS-6 foci near centrioles using 3D-STORM revealed a range of 60-150 nm [30]. Comparably, an average of 276 molecules of SAS-6 have been estimated to be present in the vicinity of centrioles [12]. In both cases, it is likely that at least some fraction of the this SAS-6 pool is integrated into cartwheels. Along this line of thought, a recent paper could not detect cenriole assembly defects in a SAS-6 degradation system despite a significant reduction in its levels [14]. Under the light of our theoretical results, cartwheels are expected to be shorter in this system and it would be interesting to test this expectation by measuring their length..

Testing this model requires correlating SAS-6 levels with cartwheel numbers and length. Ideally, this should be done at a single-cell level in a synchronised population in G2, i.e. after centriole duplication. At a finer scale, it requires a readout of cartwheel assembly and stacking kinetics. This is challenging because human cartwheels are extremely small structures. Recent studies have been successful in studying cartwheel assembly and stacking *in vitro*. First, several authors have produced and measured cartwheel-like stacks *in vitro* [14, 26]. Second, a specialised atomic-force microscopy protocol has enabled cartwheel ring assembly to be imaged live [27]. In principle, it should be possible to manipulate the concentration of SAS-6 and study how it affects the number of stacks or their length. These systems may be used to study basic properties of cartwheel assembly, as modelled here, bearing in mind that they do not reflect *in vivo* centriole duplication. Experimental validation of the model lies beyond the scope of this paper but should be addressed in the future.

Although the basic properties of SAS-6 self-oligomerisation have been well established in the literature it is still unclear how rings stack up. It has been suggested that rings are first assembled and then loaded onto a growing cartwheel. Such stacking mechanism is distinct from the we assumed and understanding its consequences for centriole number control would demand specific modelling.

### Generality and relation to other models

A model of cartwheel assembly similar to ours has been recently employed to estimate reaction rates associated SAS-6 self-assembly and dissociation [36]. This model has not been interpreted in the context of centriole number control, let alone the possibility that this is enforced by the unique stacking mechanism we propose. Other models have focused on the issue of copy number control postulating mechanisms that require a spatially explicit components. In contrast, our model relies only on the existence of multiple intermediates in the cartwheel assembly pathway, which can be sequestered by the elongation of the preformed cartwheels. The sequestering of the building block intermediates represents the inhibitory mechanism that proceeds without demanding the physical existence of a diffusive inhibitor like in lateral inhibition. Since mutants of this diffusive inhibitor will disrupt copy number control in the models of auto-activation and lateral inhibition, we expect our model to be more robust.

Instead, our model is akin to kinetic proofreading. Kinetic proofreading was originally proposed to explain the low error rates observed in DNA replication and protein translation [37], stipulating that errors (i.e. wrong reaction products) would be minimised by a correction mechanism that iterates multiple times over a given substrate. These ideas were later co-opted to explain the narrow specificity of T-cell receptors [38, 39]. In contrast with the latter views, in which TCR signalling is made specific because a multi-iterative process can only be completed in a narrow range of conditions, we take advantage of the complementary broad range of conditions in which such a process fails, preventing extra cartwheels from arising.

Ultimately, how centrioles duplicate exactly once per cell cycle remains an unresolved question in cell biology. Our models predict that formation of one and only one cartwheel should be robust to random fluctuations in SAS-6 expression within a cell population. A unique prediction of our model is that such stochasticity should be reflected in cartwheel length variation, which is a property that has been hitherto overlooked. and that both cartwheel numbers and length are increase monotonically with SAS-6 levels. Testing these predictions can help further our understanding of how centriole numbers are regulated in proliferating cells. Here, we propose a simple and original mechanism for copy number control that is reminiscent of basic biochemical principles and establishes quantitative relationships between stoichiometry, number, and length.

## Models and Methods

In this section we present detailed formulation of the models used in this study and describe the simulation methods.

### SAS-6 dimer fate model

We assume a large enough volume around a single pre-existing centriole that is large enough to allow for mass action kinetics. FCS data suggest that homodimers are the most abundant SAS-6 species in the cytoplasm [30]. Thus, we consider in our models that SAS-6 dimers are the fundamental cartwheel building block. We consider a single SAS-6 dimer in the presence of a pre-existing cartwheel. We assume that this dimer can progress sequentially through *i* intermediate stages, until forming a ring of size *r*, thus establishing a new cartwheel. Alternatively, intermediates of size *i* can be diverted by stacking up on top of the pre-existing cartwheel. This system can be represented by the following chemical equations:

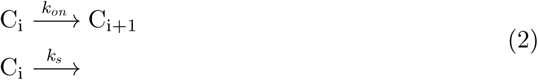

where C_i_ denotes an intermediate containing *i* dimers, with *i< r*. We assume that the addition of other dimers to the growing intermediate occurs at a constant rate *k*_*on*_, and that stacking of the intermediate on a pre-existing cartwheel occurs at a rate *k*_*s*_. The process stops when the final product *C*_*r*_ is assembled.

Let *T*_*i*_ denote the residence time of the intermediate containing *i* dimers. We assume all *T*_*i*_ variables are independent and identically distributed, following an exponential distribution with rate parameter (*k*_*on*_ + *k*_*s*_). We define *O*_*i*_ as the sum of residence times *T*_1_,…,*T*_*i*_. The variable *O*_*i*_ is correctly interpreted as the time it takes for completing *i* reactions. The the cumulative distribution of *O*_*i*_ is given by the convolution of the cumulative distributions of *T*_1_,…,*T*_*i*_. Thus, it can be defined recursively as:

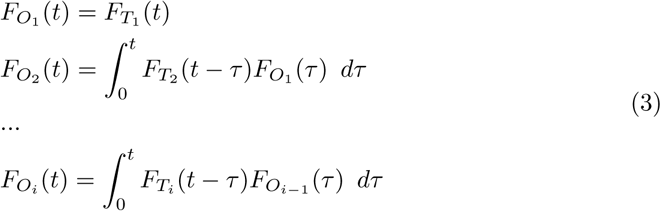

It is worth noting that these expressions imply that the initial dimer completed at least *i −* 1 reaction steps leading to larger intermediates. In other words, it requires that the initial dimer was not diverted to the elongating chartwheel up to time *t*. Thus, it is convenient to weight Eq. 3 by the expected value of the probability that the dimer integrated an *i −* 1 intermediate up to time *t*. This yields:

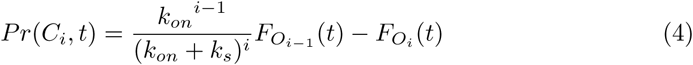

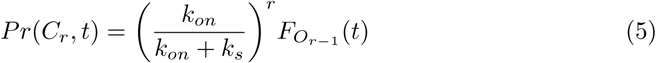

where equation Eq. 4 represents the probability distribution that the dimer is in step *i* of the assembly pathway, at time *t*, and Eq. 5 is the probability distribution that the dimer seeded a new cartwheel at time *t*.

Finally, taking the limit of Eq. 5 when *t →∞* we obtain:

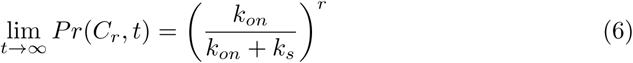

which is the maximal probability that the initial dimer gives rise to an extra cartwheel. This is minimal condition that allows for supernumerary centrioles to be formed.

### Stochastic cartwheel assembly model

We assume a constant influx rate σ of SAS-6 dimers, as a simplification of all cellular processes related to SAS-6 expression, degradation, and recruitment. SAS-6 dimers are known to form higher-order assemblies via each of their two N-termini. We describe oligomerisation as a second-order process involving any two intermediates containing *I* and *j* dimers, subject to *i* + *j* ≤*r*. In other words, oligomers can combine to produce any structure that is not larger than a complete ring. A complete ring is an oligomer with *r* dimers. It should be noted that ring formation requires *r* dimers to be bound and an additional reaction to close the ring. For simplicity, we assume this reaction is fast enough that it can be neglected. In addition, we assume that *r* dimers are irreversibly bound. Thus, a ring effectively gives rise to a new cartwheel. Furthermore, we assume that all oligomerisation reactions, including the ones yielding complete rings, occur at a rate *k*_*on*_. Note that the actual oligomerisation reactions include the two exposed N-termini in a given SAS-6 oligomer. Intermediates containing *k< r* dimers are assumed to dissociate into any of their constituents, containing *i* and *j* dimers (*k* = *i* + *j*), with a rate *k*_*off*_.

In practice, we envision stacking as identical to ring assembly atop a pre-existing cartwheel. This requires vertical interactions between the oligomer and the stack, as well as lateral interactions with other molecules in the top layer of the cartwheel. For simplicity, we assume that this is achieved in a single reaction step, with a rate *k*_*s*_. We assume free-diffusing oligomers containing *i* dimers can bind to the *j* dimers in the top layer of the cartwheel, at a rate *k*_*s*_. If *i* + *j* = *r*, a new ring is completed on the stack. As is the case with ring formation, stacking reactions are assumed to be irreversible. Technically, our model makes no distinction on the directionality of stacking; we refer to “top” merely as a verbal device for ease of comprehension. Nevertheless, it should be recalled that in some cellular systems cartwheel elongation seems to be proceed unidirectionally from the bottom [40].

This system can be represented by the following chemical equations:

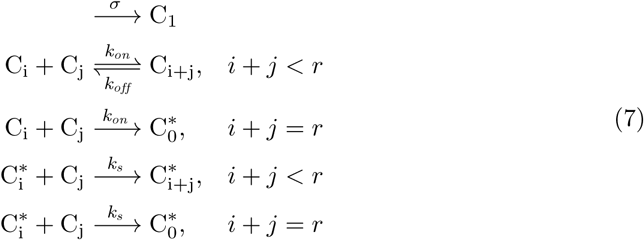

using the previous notation and * to specify stacks. The first equation corresponds to SAS-6 influx; the second equation indicates bimolecular oligomerisation and dissociation; the third equation denotes irreversible assembly of complete rings; the fourth equation represents stacking by combination of a cartwheel with an incomplete top layer (an incomplete stack); the fifth equation represents addition of a complete ring to a stack by adding *r* dimers to its top layer. Note that the species 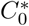 replaces *C*_*r*_ and includes all stacks whose top layer is a complete ring. Thus, cartwheels of different length are not formally represented in this system. However, they can still be tracked in the simulations by considering stacks of different lengths as distinct chemical species.

On a final note, we assumed oligomerisation and stacking is purely arithmetic. In other words, we ignore any steric hindrance between interacting SAS-6 structures. This issue cannot be easily addressed in our discrete stochastic framework but we acknowledge it may be particularly relevant for stacking.

### Variants and alternative models

In the previous model we assumed that ring completion and stacking are irreversibly. We relaxed these assumptions to assess their impact on cartwheel formation and elongation. Alternatively, we assumed that complete rings can also dissociate at a constant rate *k*_*off*_ and that oligomers can dissociate from the top layer of a stack at a rate *k*_*u*_. Thus, ring disassembly and stack dissociation reactions are considered to be the backward reactions of ring completion through oligomerisation and stacking, respectively (third and fifth equations in system Eq. 7.

In the limited stacking model, we set a maximum cartwheel length *h*, which can take positive integer values. In the simulations, we assume that *k*_*s*_ = 0 for cartwheels of length *h*.

Regarding the feedback model, we assume that there is an arbitrary number of components that are activated sequentially. If the time for activation of the next component is exponentially distributed, then the time to complete the entire feedback sequence is the sum of each activation time, which is gamma-distributed with shape parameter α and rate ρ. In our simulations, we set a flag that is checked once the first cartwheel is formed and triggers the feedback loop. In practice, this involves drawing a random number from a gamma distribution with shape parameter α and ρ, after which the simulations are stopped.

### Analysis and simulations

We obtained analytical solutions for the probability distributions of Eq. 4 and Eq. 5 at time *t* using built-in functions in Mathematica. The more realistic models were implemented and simulated using Python, using the Gillespie algorithm. No SAS-6 species are present initially for all simulations. Parameter values are indicated where appropriate; by default σ = *k*_*on*_ = *k*_*off*_ = *k*_*s*_ = 1 and *r* = 9. Unless stated otherwise, we performed 1000 simulations of the indicated model(s), in the time-window = *t*[0, 100] starting with no SAS-6 molecules present.

## Acknowledgments

We are grateful to all members of the Quantitative Organism Biology and Cell Cycle Regulation labs for insightful discussions and to Carla A. M. Lopes, Catarina Nabais, Catarina Peneda and Sonia Gomes Pereira, in particular, for reading and commenting an earlier version of the manuscript.

---

**Table S1.** Parameters across all models.

| Parameter | Definition | Units | Range |
| --- | --- | --- | --- |
| $\sigma$ | SAS-6 influx rate. Simplification of SAS-6 expression and degradation dynamics, as well as centrosomal recruitment. | molecules $t^{-1}$ | $[0, \infty[$ |
| $k_{on}$ | Oligomerisation rate. Rate at which two SAS-6 oligomers of the same or different sizes combine. | molecules<sup>-1</sup>$t^{-1}$ or molecules $t^{-1}$ | $[0, \infty[$ |
| $k_{off}$ | Dissociation rate. Rate at which a SAS-6 oligomer dissociates into any combination of two of its constituents. | molecules $t^{-1}$ | $[0, \infty[$ |
| $k_s$ | Stacking rate. Rate at which a SAS-6 oligomer combines with a cartwheel by stacking on top of it. | molecules<sup>-1</sup>$t^{-1}$ | $[0, \infty[$ |
| $r$ | Ring size. Number of SAS-6 oligomers in a complete ring, necessary to form a new, distinct cartwheel. | molecules | $\mathbb{N}$ |
| $h$ | Maximum cartwheel length. Maximum number of stacked rings in a cartwheel, at which further stacking reactions are blocked. | rings | $\mathbb{N}$ |
| $\rho$ | Mean feedback time. Mean time after the first cartwheel formation event at which further synthesis and elongation are prevented (i.e. the simulation is stopped). | $t$ | $[0, \infty[$ |
| $\alpha$ | Shape parameter of the gamma-distribution of feedback times. When $\alpha=1$ feedback time simplifies to an exponential distribution. | Adimensional | $[1, \infty[$ |

**Fig. S1.**
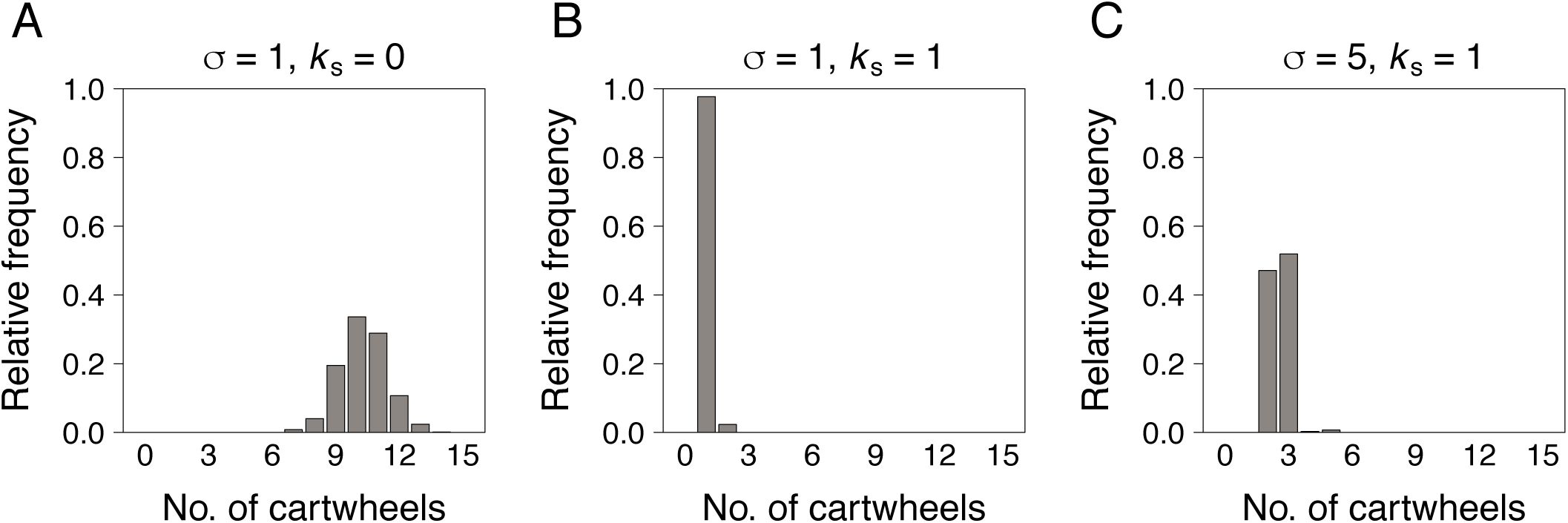
Final cartwheel number distributions. (A-C) Relative frequency of simulations with the indicated number of cartwheels, at the stopping time. We performed 1000 simulations with no SAS-6 molecules initially present and stopped them at time *t* = 100; time *t* is measured in arbitrary units (a.u.). For all simulations, *k*_*on*_ = *k*_*off*_ = 1. Note that *k*_*s*_ =0 represents absence of stacking. See also Methods section for default initial and stopping conditions, and parameter values for the simulations.

**Fig. S2.**
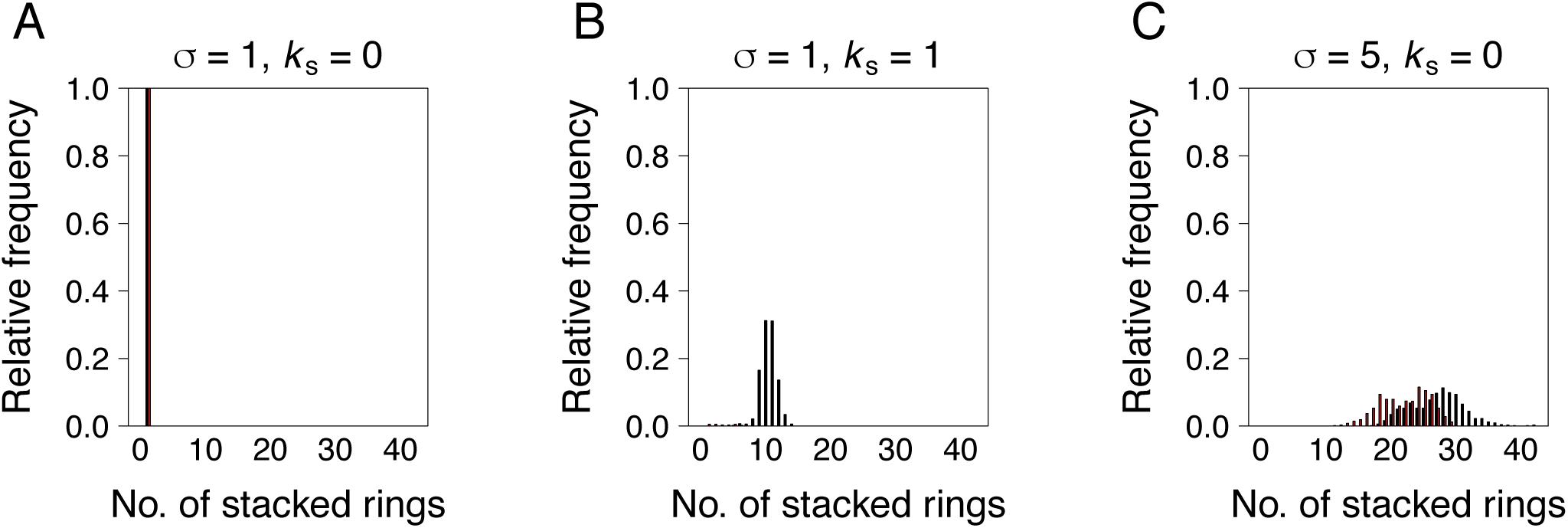
Final cartwheel length distributions. (A-C) Relative frequency of simulations where the first (black) and second (red) cartwheels contained the indicated number of stacked rings, at the stopping time. We used default simulation settings as indicated in Figure S1 and in the Methods section.

**Fig. S3.**
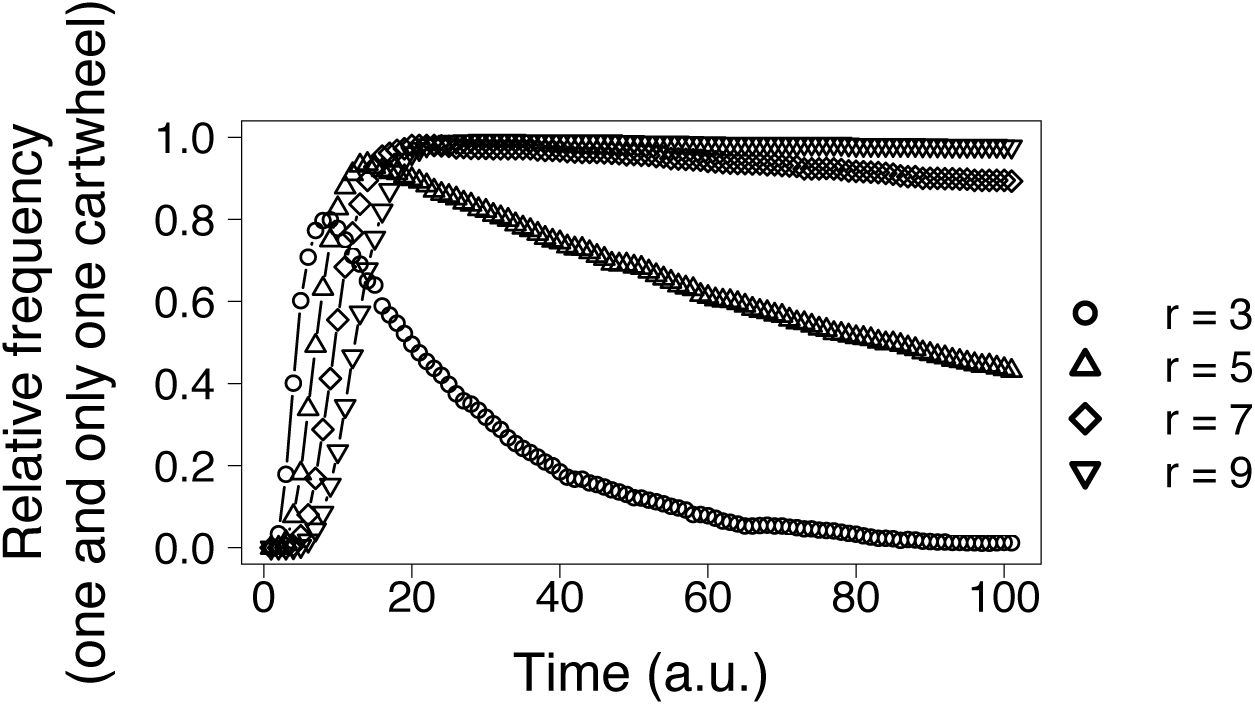
The probability of forming one and only one cartwheel increases with ring size. Relative frequency of simulations containing one and only one cartwheel as a function of time, for the indicated values of *r*. We used default simulation settings as indicated in Figure S1 and in the Methods section.

**Fig. S4.**
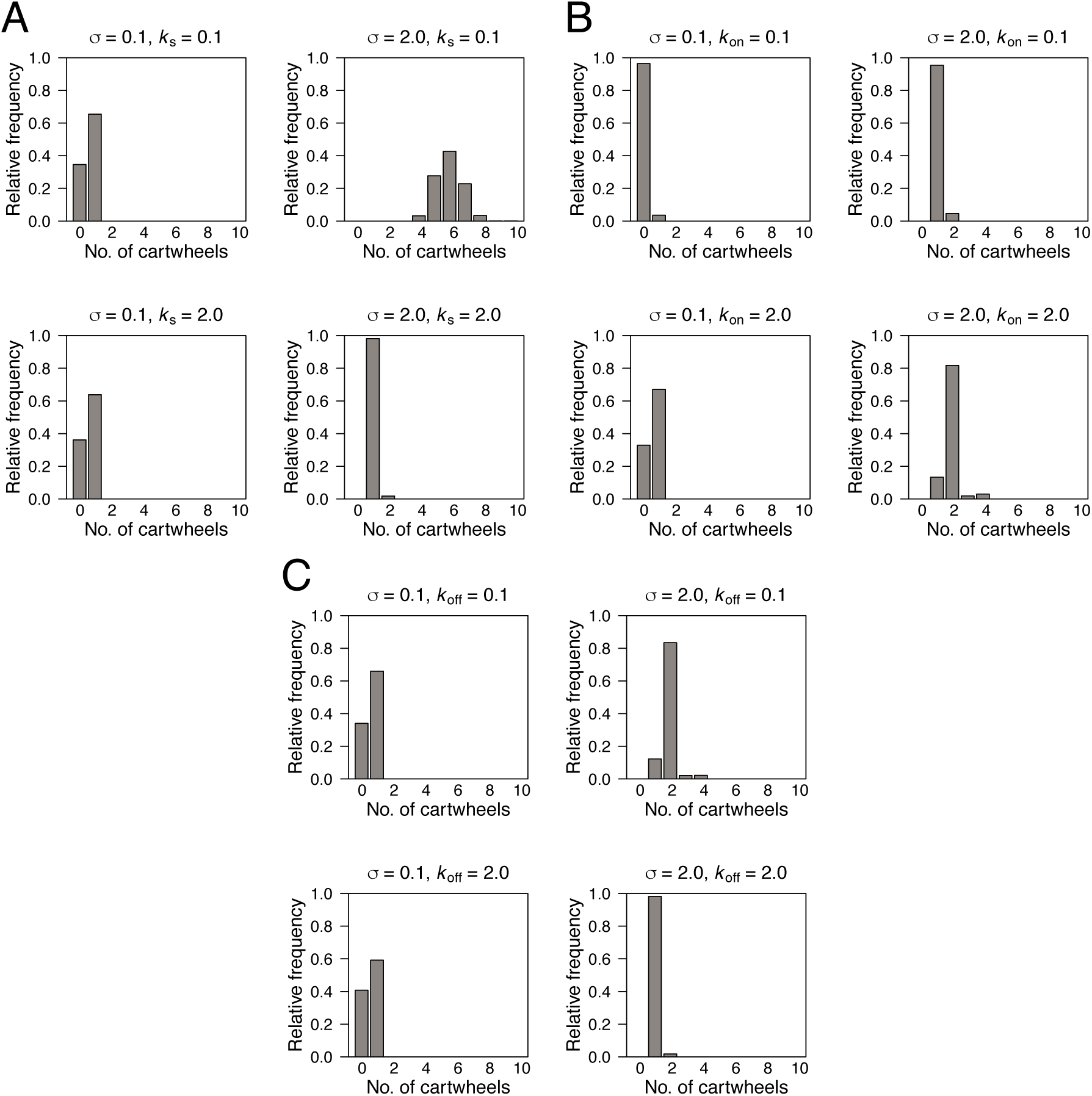
{(A-C) Final distribution of cartwheel numbers as a function of influx and different reaction rate parameters. We used default simulation settings as indicated in Figure S1 and in the Methods section.

**Fig. S5.**
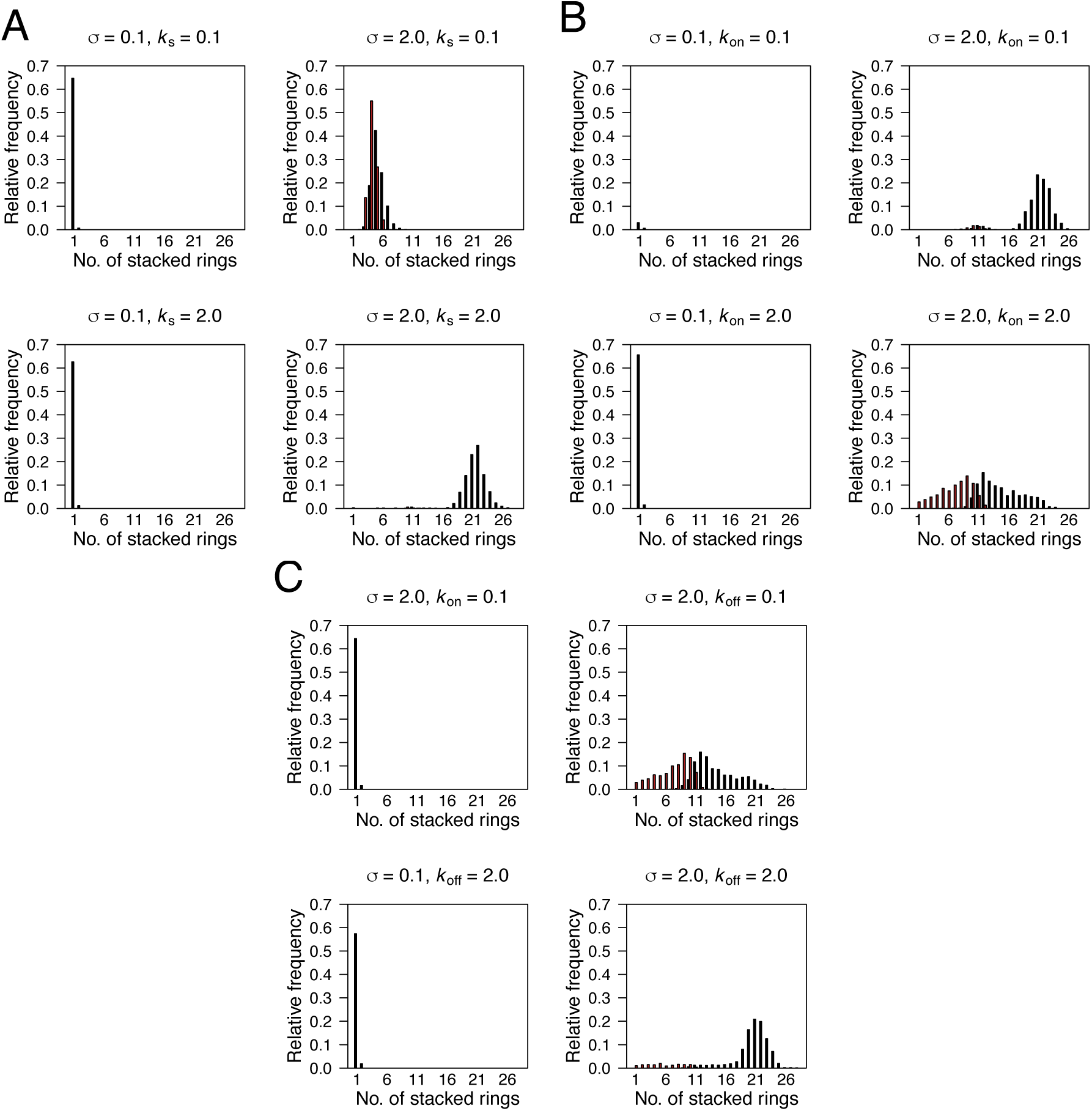
A-C) Final distribution of cartwheel lengths as a function of influx and different reaction rate parameters. We used default simulation settings as indicated in Figure S1 and in the Methods section.

**Fig. S6.**
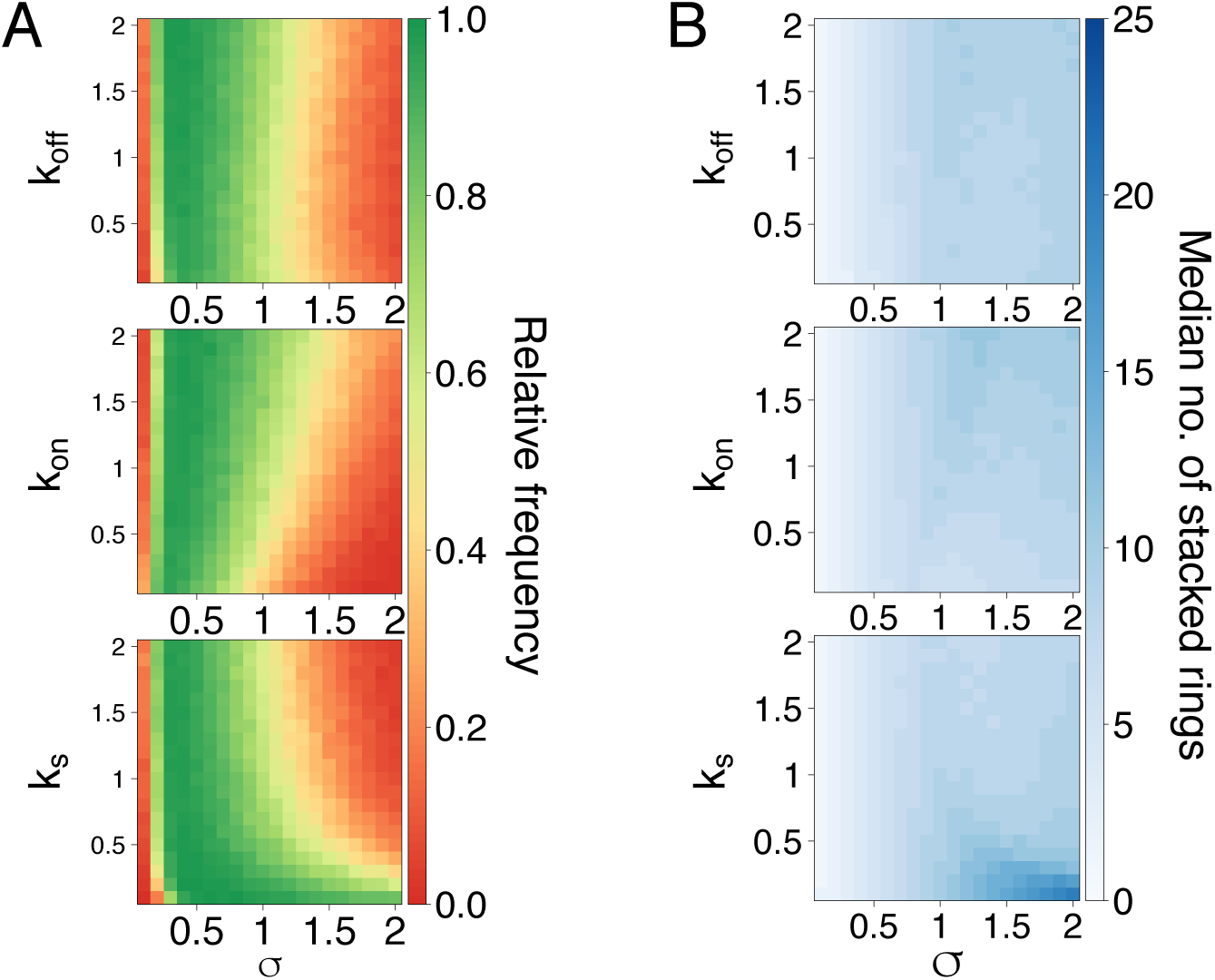
Probability of forming one and only one cartwheel as a function of influx and reaction parameters, assuming reversible ring formation and stacking. Relative frequency of simulations containing a single cartwheel at time *t* = 100, under the model assuming reversible ring assembly and stacking. We varied the parameter values indicated in the x-and y-axes in steps of 0.1, yielding a total of 400 pairwise combinations of parameter values per plot. We used default simulation settings as indicated in Figure S1 and in the Methods section, and set *k*_*u*_ = 1. Note that *k*_*s*_ =0 represents absence of stacking.

**Fig. S7.**
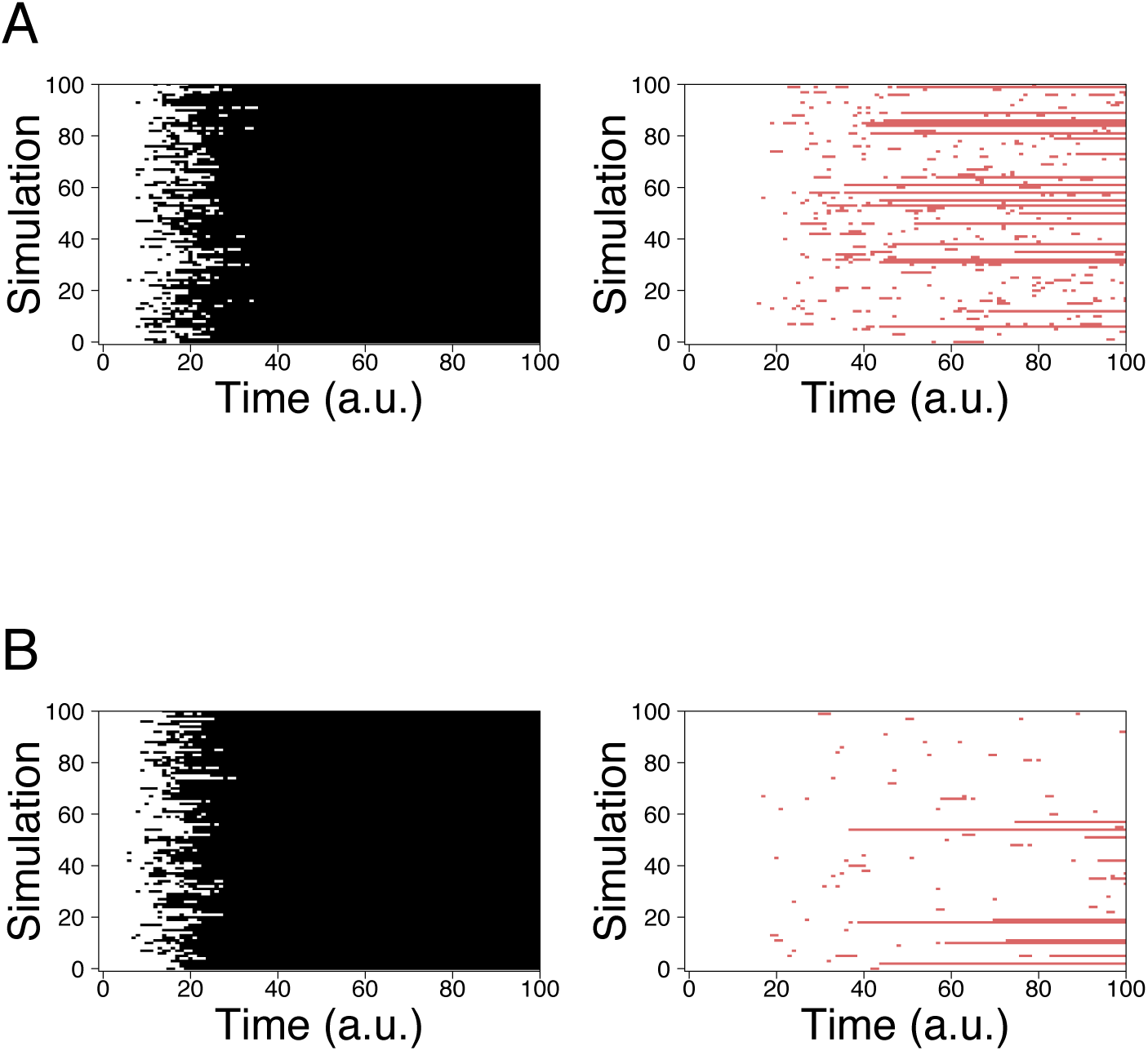
The first assembled cartwheel is stably maintained whereas the second one is dynamically assembled and disassembled. Presence of the first (black) and second (red) cartwheels along time, under the model assuming reversible ring assembly and stacking, in a random sub-sample of 100 out of 1000 simulations. We used default simulation settings as indicated in Figure S1 and in the Methods section, and set *k*_*u*_ = 1. A - *k*_*s*_ = 1; B - *k*_*s*_ =5

**Fig. S8.**
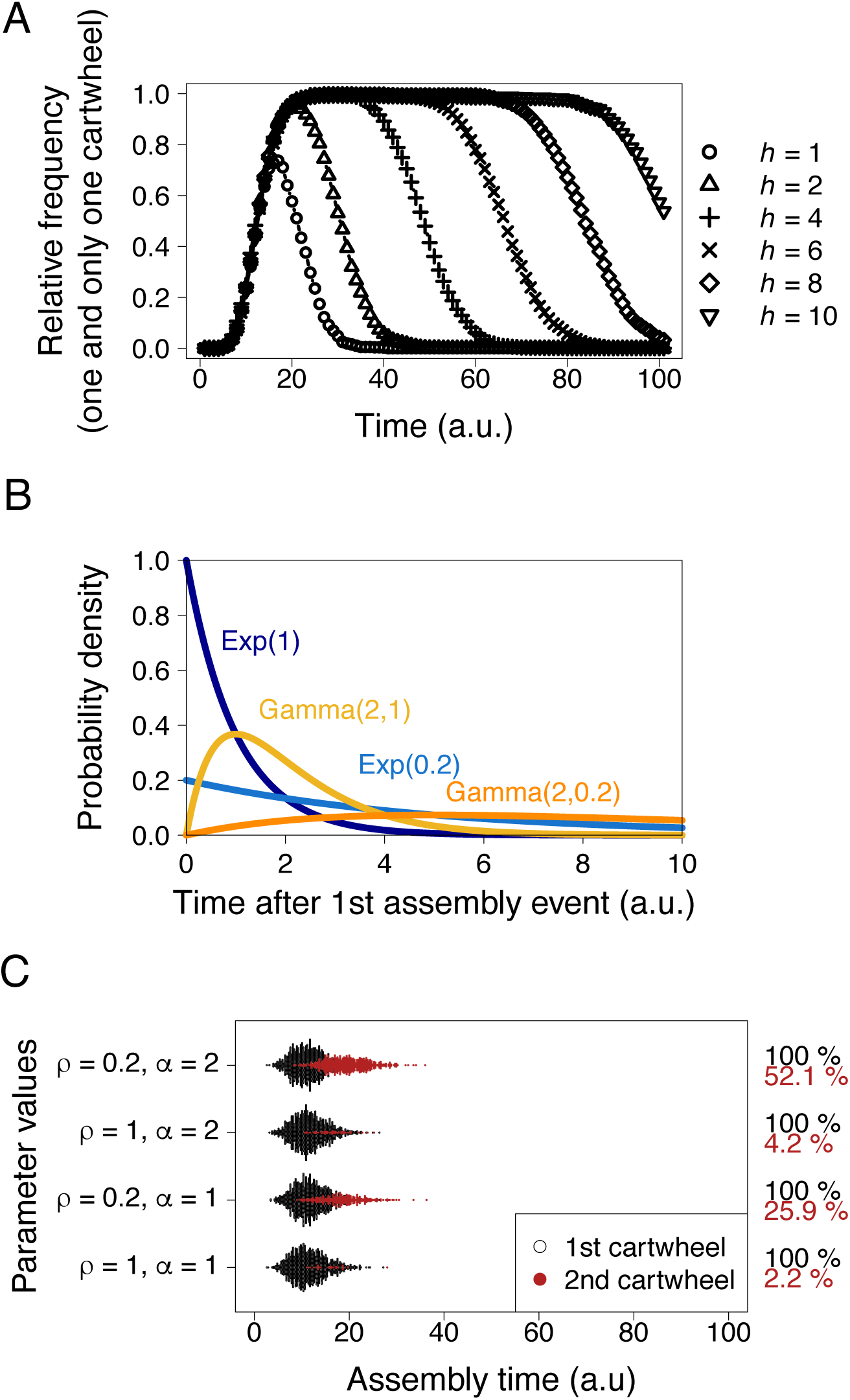
Additional features of the limited stacking and feedback models. A - Assembly of supernumerary cartwheels is progressively delayed as the maximum cartwheel length *h* increases. We used default simulation settings as indicated in Figure S1 and in the Methods section. B - The distribution of feedback times simplifies to an exponential distribution for *α* = 1, with mean ρ. Otherwise, it is gamma-distributed. C - Assembly times of the second cartwheel as a function of feedback parameters. We used default simulation settings as indicated in the Methods section.

